# Gene editing of the multi-copy H2A.B gene family by a single pair of TALENS

**DOI:** 10.1101/233379

**Authors:** Nur Diana Anuar, Matt Field, Sebastian Kurscheid, Lei Zhang, Edward Rebar, Philip Gregory, Josephine Bowles, Peter Koopman, David J. Tremethick, Tatiana Soboleva

## Abstract

In view of the controversy related to the generation of off-target mutations by gene editing approaches, we tested the specificity of TALENs by disrupting a multi-copy gene family using only one pair of TALENS. We show here that TALENS do display a high level of specificity by simultaneously knocking out the function of the three genes that encode for H2A.B.3. This represents the first described knockout of this histone variant.

---

Despite being discovered over a decade ago ^1^, the *in vivo* importance of the mammalian histone H2A variant H2A.B remains unknown. H2A.B appeared late in evolution in mice (H2A.B.3) and humans (H2A.B), and it is predominantly expressed in the testis with low expression levels in the brain ^2,3^. The availability of a H2A.B.3 knockout would be an invaluable tool for investigating the importance of H2A.B, however, as the H2A.B.3 protein is expressed from three different genes, this is a technically challenging feat. Our aim was to devise the most efficient and specific strategy to inhibit the function of all three genes. Here, we report a simplified TALEN approach that achieves this goal.

Previously, transcription activator-like effector nucleases (TALENs) were the choice to perform gene and genome editing ^4,5^ but more recently this technology has been superseded by the CRISPR-Cas9 system ^6^. However, while improvements are continually being made ^7-9^, one issue has been the extent of off-targets mutants generated with the use of CRISPR-Cas9, which appears to be greater when compared with TALENs ^10-13^. One attractive feature of TALENs, which enables a higher level of specificity, is that the FOK1 nuclease domain will only cleave DNA when dimerized, which occurs when the two TALE-domains bind to DNA (on opposite strands) in close proximity to each other. Here, we used TALENs to genetically impair the function of all three H2A.B.3 encoding genes and moreover, we used only one pair of TALENS to do this. To our knowledge, this has not been done before.

There are three H2A.B.3-encoding genes (H2Afb3, Gm14920, H2Afb2, which are >92% identical) plus a pseudogene (Gm14904) all located on the X chromosome in the mouse (this pseudogene was present in the Ensemble release 57 but subsequently removed) (**Supplementary Table 1**). We confirmed the expression of the three H2A.B.3-encoding genes in the testis and as expected, the pseudogene was not expressed (**Supplementary Fig 1**). In order to test the specificity of TALENs against the H2A.B.3 gene family, two groups of H2A.B.3-targeting TALENs were designed. The first group of 12 TALENs were designed to target only one gene, H2Afb3. The second group of 7 TALENs was design to target all three H2A.B.3 genes (**Supplementary Table 2**).

The activity and specificity of the TALENs were tested by employing a Dual-Luciferase Single Strand Annealing Assay (DLSSA) and by the Cel1 cleavage assay. The TALEN plasmids were co-transfected into Neuro2a cells in pairs in various combinations (**Supplementary Fig. 2**). The results for the first group showed that all TALEN pairs, in different combinations, had the highest activity for the H2Afb3 gene, with some TALEN pairs indeed only cleaving the H2Afb3 gene (e.g. 101408:101412). The second group of TALENs showed nuclease activity for all genes, with 2 pairs (101421:101422 and 101421:101423) having the highest activity. Cel1 assays confirmed these findings (**Supplementary Fig. 3**). These *in vitro* results demonstrate the high specificity of TALENs; some of the targeted DNA sequences of the first group, designed against the H2Afb3 gene, differed by only 2-3 base pairs compared to the second group, and yet specificity for only the H2Afb3 gene was achieved (**Supplementary Fig. 4**). The TALEN pair 101421:101422 was used to knock out (KO) the function of all three H2A.B.3 genes in mice (**Supplementary Fig. 4**).

Prior to creating the H2A.B.33 KO, we determined whether any off-target genomic locations, which have limited sequence homology to the TALEN pair 101421:101422, can be identified by using NCBI BLAST against the mouse reference genome mm10 built on the C57BL/6J strain. Even relaxing this BLAST search to only 8 bp out of the 15 bp DNA recognition sequence returned no matches (data not shown).

In order to produce H2A.B.3 KO mice, FVB/NJArc mice (derived in Jackson Laboratories and sourced from Australian Animal Resource Centre (ARC), hence an extra suffix, JArc) were used. Capped, poly-adenylated and purified *in vitro* transcribed mRNA was adjusted to 10 ng/μ! for each TALEN pair (101421:101422) and injected into 532 fertilised one-cell embryos and cultured overnight to the 2-cell stage. 153 surviving 2-cell embryos that displayed normal morphology were implanted in pseudo pregnant females, resulting in the birth of 19 pups. Of these 19 pups, 9 contained TALEN-induced H2A.B.3 mutations. Most interestingly, TALENs activity produced an all-or-nothing effect, i.e., for each of the 9 founder pups, all three H2A.B.3 genes were mutated while for the 10 unmodified pups, no gene copies of H2A.B.3 were modified (Table 1). Moreover, in female pups, both alleles of H2Afb3, Gm14920 and H2Afb2 were modified, and each H2A.B.3 gene contained a unique indel as the result of TALEN activity (identified by Sanger sequencing). Therefore, one pair of TALENS can edit 6 homologous alleles simultaneously.

**Table 1.** Genotype of all 19 pups born following TALEN injections. Wild type (Wt.) offspring. Δn,deletion of n nucleotides; +n, insertion of n nucleotides. * denotes mice that were used as founders to establish H2A.B.3KO colonies.

| # | Pup ID | Genotype by Sanger sequencing |  |  |
| --- | --- | --- | --- | --- |
|  |  | H2Afb3 | Gm14920 | H2Afb2 |
| 1 | ♀ L74-1 | XX (wt) | XX (wt) | XX (wt) |
| 2 | ♀ L74-2 | XX (wt) | XX (wt) | XX (wt) |
| 3 | ♀ L74-3 | XX (wt) | XX (wt) | XX (wt) |
| 4 | ♂ L74-4 | XY (wt) | XY (wt) | XY (wt) |
| 5 | ♂ L74-5* | $X^{\Delta 5 +286}Y$ | $X^{\Delta 15}Y$ | $X^{\Delta 17}Y$ |
| 6 | ♀ L79-1 | $X^{\Delta 10}X^{\Delta 10}$ | $X^{\Delta 12}X^{\Delta 12}$<br>Chimera of b/$\Delta$/c | $X^{\Delta 15}X^{\Delta 15}$<br>Chimera of c/$\Delta$/b |
| 7 | ♀ L89-1* | $X^{\Delta 22}X^{\Delta 160}$ | $X^{\Delta 5}X^{\Delta 17}$<br>Chimera c/$\Delta$/b) | $X^{\Delta 5}X^{\Delta 5}$ |
| 8 | ♂ L89-2 | $X^{\Delta 65}Y$ | $X^{C/T, A/G}Y$ | $X^{\Delta 42}Y$<br>Chimera b/$\Delta$/c) |
| 9 | ♂ L89-3 | $X^{\Delta 26}Y$<br>Chimera c/$\Delta$/a) | $X^{\Delta 10}Y$<br>Chimera b/$\Delta$/c) | $X^{\Delta 10}Y$<br>Chimera b/$\Delta$/c) |
| 10 | ♀ L90-1 | XX (wt) | XX (wt) | XX (wt) |
| 11 | ♀ L90-2 | XX (wt) | XX (wt) | XX (wt) |
| 12 | ♀ L90-3* | $X^{\Delta 28}X^{\Delta 5}$ | $X^{\Delta 10}X^{\Delta 10}$ | $X^{\Delta 64}X^{\Delta 64}$ |
| 13 | ♀ L90-4 | $X^{\Delta 15}X^{\Delta 10}$ | No successful PCR amplification | No successful PCR amplification. |
| 14 | ♀ L90-5 | XX (wt) | XX (wt) | XX (wt) |
| 15 | ♀ L90-6 | $X^{\Delta 17}X^{\Delta 17}$<br>Chimera c/$\Delta$/a) | $X^{\Delta 5}X^{\Delta 5}$<br>Chimera b/$\Delta$/c) | $X^{\Delta 5}X^{\Delta 5}$<br>Chimera c/b/$\Delta$/c) |
| 16 | ♀ L90-7 | XX (wt) | XX (wt) | XX (wt) |
| 17 | ♂ L90-8 | $X^{\Delta 16}Y$ | $X^{\Delta 17}Y$<br>Chimera c/$\Delta$/c/b) | No successful PCR amplification. |
| 18 | ♂ L90-9 | XY (wt) | XY (wt) | XY (wt) |
| 19 | ♂ L90-10 | XY (wt) | XY (wt) | XY (wt) |

Most of the mutants carried small deletions. Interestingly, in 2 founders, L90-4 and L90-8, at least one of the H2A.B.3 genes (usually gm14920 or H2Afb2) was not amplifiable by PCR. However, when we used mixed primer pairs for PCR amplification (e.g. gm14920-forward and H2Afb2-reverse) we were able to detect an amplification product and following Sanger DNA sequencing, demonstrated that a fusion between gm14920 and H2Afb2 had occurred (data not shown). Also unexpectedly, when the founder L74-5 mouse, which only displayed the deletions and insertions described in Table 1, was bred with a wild-type (wt) female, the resulting G1 progeny displayed a chimera between the gm14920 and H2Afb2 genes **(Supplementary Fig. 5).** Continued breeding from these chimeric G1 H2A.B.3^-/x^ females with wt males (up to 4 generations), revealed that these additional mutations were lost after G2, but maintained original founder mutations (H2Afb3 X^Δ+286^, gm14290 X^Δ15^, H2Afb2 X^Δ17^). This can be explained by the fact that TALEN activity can be retained for several cell divisions following their injection into the 2-cell stage, thus producing a mosaic genotype, as previously observed ^14,15^. Collectively, the G1 generation of the 9 founder H2A.B.3 genetically modified mice produced 22/78 mice displaying mosaicism, but by G3 a pure non-mosaic mouse colony was derived.

The strategy for the identification of possible TALEN-induced off-target mutations in H2A.B.3^-/y^ mice was based on the sequencing of exomes of three related non-mosaic H2A.B.3^-/y^ mice from three consecutive generations (G1-G3). This strategy avoids the problem observed in previous studies where unrelated mice were compared to assess offtarget effects ^12^. Each exome sample was sequenced using 100bp paired-end reads on the Illumina HiSeq 2000 sequencer to a depth of 350 - 600x coverage, yielding between 129×10^6^ and 228×10^6^ reads per sample (**Supplementary Table 3**). The breeding began by crossing ♀L90-3 with ♂L74-5 to produce phenotype NM4-G1 H2Afb3 (X^Δ5^Y), gm14920 (X^Δ10^Y) and H2Afb2 (X^Δ64^Y) mice. Subsequent crosses between genetically modified H2A.B.3 female mice and wt males produced NM4-G2 and G3 H2Afb3 (X^Δ5^Y), gm14920 (X^Δ10^Y) and H2Afb2 (X^Δ64^Y) mice for exon sequencing. By crossing mutant females with wt. males any true off-target mutations would be identified because they would be diluted by 50% after each generation. On the other hand, natural sequence variations due to mouse strain differences would not be diluted by breeding (see below). Sequenced mouse exomes were run through our inhouse variant detection pipeline (see Online Methods) to detect SNVs, small indels and larger structural variants. Importantly, TALENS do not usually produce SNVs, thus this type of variant serves as an internal measure for mouse strain differences.

As expected, alignment with the mm10 genome identified a large number of SNVs (~1.7×10^6^ for G1 and G2, and ~0.69×10×^6^ for G3) and indels (~1.7×10^5^ for G1 and G2, ~0.6×10^5^ for G3) (**Supplementary Table 4**), however neither followed a dilution pattern but rather correlated with the original number of sequencing reads. To remove the effect of mouse strain differences, a filter for FVB/NJ was included into the alignment, which indeed greatly reduced the number of SNVs and indels (**Supplementary Table 5**). Again, the remaining SNVs and putative indels did not follow a dilution pattern following successive generations.

An additional analysis was conducted to separate heterozygous from homozygous indels ^16^. As shown (**Supplementary Table 6**), after filtering against the FVB/NJ genome, 417 NM4-G1, 420 NM4-G2 and 325 NM4-G3 heterozygous (0/1+0/2+1/2) indels remained after alignment with the mm10 genome, but again did not display a 50% dilution pattern from G1-G3. To provide further evidence that these remaining indels are due to strain differences between FVB/NJArc and FVB/NJ mice, we interrogated the 19 indels common to all three NM4 G1-G3 mice (**Supplementary Fig. 6**, the location of these putative indels are shown in **Supplementary Table 7**).

We amplified ~300-450 bp regions spanning all 19 indels and compared them to the wt FVB/NJArc mice. The results for all 19 regions (**Fig. 1a**) clearly show that the amplified fragments were identical in size for the H2A.B.3^-/y^ and wt FVB/NJArc mice. Furthermore, we sequenced 5 of these putative indels, which confirmed that the H2A.B.3^-/y^ and wt mice are identical in the nucleotide sequence where the indels were predicted to be (**Fig. 1b and c**). Further, the computational prediction tools are often inaccurate, not only in predicting the size of a putative deletion but also in falsely predicting that a deletion exists (**Fig. 1c and d**). Overall, our results suggest that the predicted indels in our H2A.B.3^-/y^ are due to DNA sequence differences in our in-house FVB/NJArc mouse strain, and computational errors in predicting indels. Finally, to detect structural variants in NM4-G1, NM4-G2 and NM4-G3, analyses using Pindel and Janda were performed. In addition to the three expected deletions in H2A.B.3 genes: H2Afb3 (5 bp), gm14920 (10 bp), a larger deletion of 64 bp in H2Afb2 was also detected (**Supplementary Fig. 7**).

**Figure 1.**
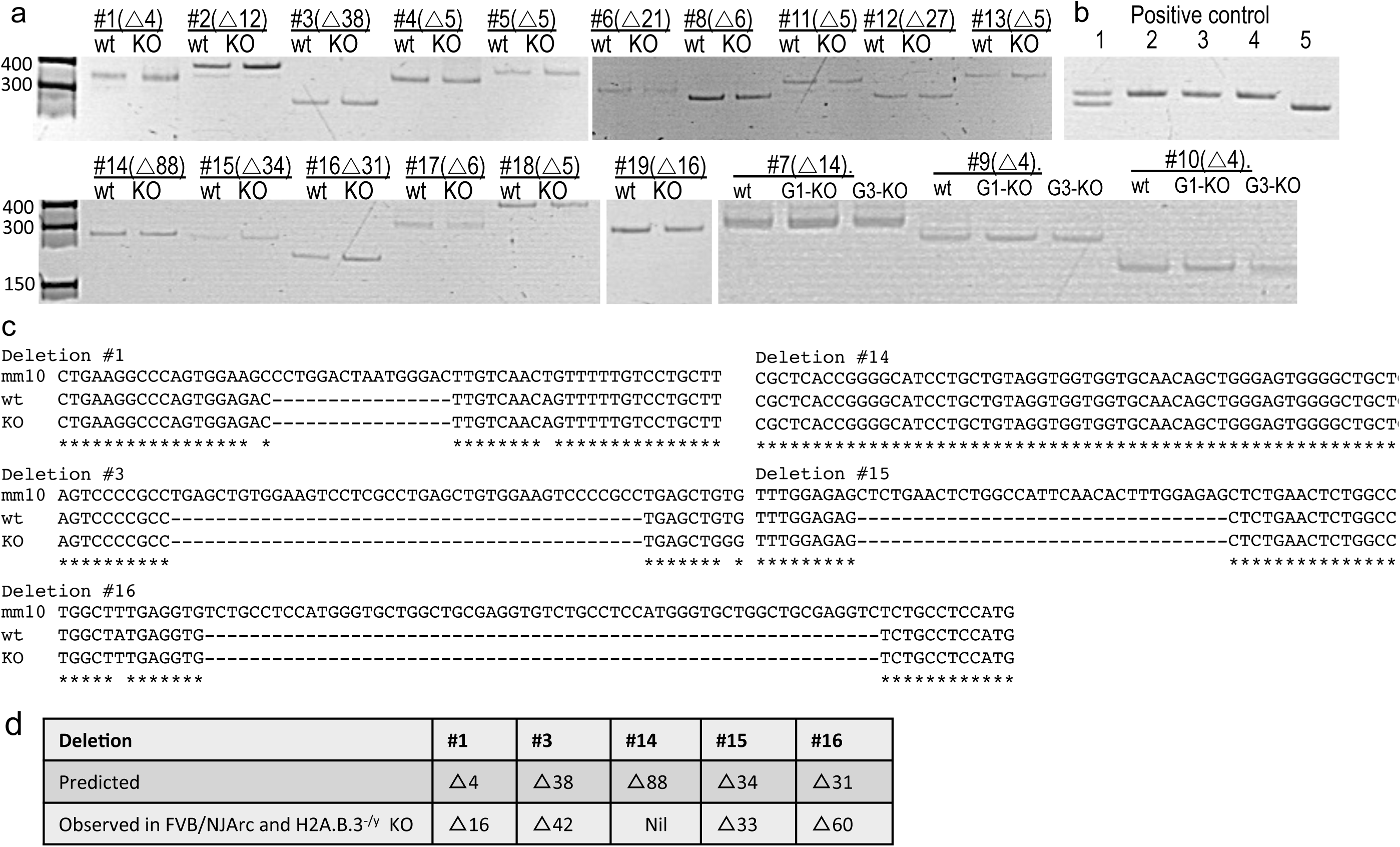
Interrogation of predicted off-target heterozygous deletions in H2A.B.3.3^-/y^ mice. (**a**) 300-400bp regions surrounding 19 common predicted heterozygous deletions were amplified from gDNA of H2A.B.3.3^-/y^ KO and FVB/NJArc in-house wt mice. PCR product sizes for wt and H2A.B.3.3^-/y^ were compared by resolving the products on a 7% polyacrylamide gel, side by side. The predicted deletion is indicated with Δ, followed by the number of nucleotides (nt) predicted to be deleted. (**b**) Positive controls showing that a deletion as small as 5-10 nt can be resolved on 7% polyacrylamide gel. Lane 1, a heterozygous Δ5nt, lanes 2, 3, 4 wt.; lane 5, homozygous Δ10nt. (**c**) 5 out of the 19 PCR products were sequenced by Sanger sequencing to show that FVB/NJArc and H2A.B.3.3^-/y^ mice have identical sequences but they are different from the mm10 reference genome. (**d**) Table showing that the prediction tools are often inaccurate at predicting the presence or the size of a putative deletion.

Despite the fact that H2A.B.3-deficient mice produce neither wt H2A.B.3 mRNA nor protein (**Supplementary Fig. 8a and b**), they can obviously reproduce. To investigate whether the loss of H2A.B.3 does affect fertility, we set up a breeding regime whereby age-matched male wt H2A.B.3 and H2A.B.3^-/y^ were mated with 3-6 month female mice and the litter sizes were recorded (**Supplementary Fig. 8c**). The results showed that H2A.B.3^-/y^ mice were subfertile, producing significantly smaller litter sizes and thus demonstrating that H2A.B.3 is required for fertility.

In conclusion, a major ongoing question is whether a RNA (e.g. CRISPR)- or protein (e.g. TALEN)-based mechanism will ultimately prove to be more specific for genome editing and this will be critical for future medical therapies. In this study, we have successfully applied the TALEN technology, whereby we used one pair of TALENs to specifically and simultaneously disrupt three gene copies of a gene family. Many human diseases are caused by gene copy number variations ^17^ and therefore the use of a limited number of genome editing enzymes might help in the future treatment of such diseases. The use of only one pair of TALENs, rather than multiple pairs, reduces the complexity of the TALEN approach (and hence the cost) and would reduce the likelihood of mosaicism and most importantly, reduce the possibility of off-target mutations. Indeed, our rigorous exome analyses did not reveal any TALEN-induced off-target mutations. We also found that the computational tools used to predict putative SNVs and indels have limitations, and therefore wet-lab validation is required before any conclusions can be made about off-target mutations.

Finally, the TALEN approach did expose several interesting phenomena. Firstly, the simultaneous knockout of highly homologous H2A.B.3 genes leads to the formation of chimeras between these genes. As every chimera contained a small deletion, it is likely that terminal microhomology, a mechanism of NHEJ, was involved ^18^. It is important to point out that other technologies that rely on NHEJ for inducing mutations, like CRISPR/Cas9 technology, will also likely produce chimeras between highly homologous genes. Several rounds of backcrossing can successfully eliminate such small-scale genomic rearrangements. Secondly, we observed an all-or-nothing effect whereby the TALENs either modified all 3 H2A.B.3 genes or modified none, which reduced the need for a more complex breeding and screening strategy to establish a H2A.B.3 KO colony.

## METHODS

Methods are available in the online version of the paper.

Note: Any Supplementary Information is available on the online version of the paper.

## ONLINE METHODS

### TALEN Design

The TALENs were assembled using the method previously described ^5,19^ The TALEN target sites were searched using a computer script developed in Sangamo. The four most common repeat variable diresidue (RVD), NI, HD, NN and NG, were used to recognize bases A, C, G and T, respectively. The NK RVD was sometimes used to specify the 3′ half repeat T base. TALENs were generated by linking several pre-made TALE repeat blocks in the forms of monomer, dimer, trimer, or tetramer. The desired repeat blocks were PCR amplified using position-specific primers. The PCR products were pooled, digested with BsaI and ligated into the TALEN expression vector, pVAXt-3Flag-NLS-TALE-Bsa3-C63-FokI, which had been linearized with the BsaI restriction enzyme. Ligations were transformed into *E. coli* competent cells, and single clones were picked and sequence verified.

### Dual-Luciferase Single Stranded Annealing Assay

The Dual-Luciferase Single Stranded Annealing Assay (DLSSA) is based on the Dual-Luciferase Reporter Assay System from Promega (Madison, WI). In the DLSSA, the Firefly luciferase reporter gene is split into two inactive fragments with overlapping repeated sequences separated by stop codons and one of the three H2A.B.3 genes. Introduction of a double strand break in the H2A.B.3 gene by the TALEN pair initiates recombination between the flanking repeats by the single stranded annealing pathway and produces an active luciferase gene. Neuro2a cells (ATCC #CCL-131) were cultured in Dulbecco’s Modified Eagle Medium (DMEM: Cellgro Cat#: 10-013-CV) plus 10% FBS and 2mM L-Glu (PSG optional antibiotic). One day before transfection, 20,000 cells per well were seeded into a 96-well plate. Cells were transiently transfected using Lipofectamine 2000 (Thermo Fisher Scientific) with four plasmids including a pair of TALENs, the SSA Firefly luciferase reporter, and the internal control Renilla Luciferase reporter (6.25 ng DNA for each component). The luciferase assay was performed 24 hours post-transfection according to manufacturer’s instructions. TALEN activity was measured as the ratio of the Firefly and the Renilla luminescence units.

### Surveyor Nuclease Assay

The assay has been described in detail previously ^5^. Briefly, mouse Neuro2A cells (2 x 10^5^) were seeded into a 96-well plate the day before transfection. The cells were then transfected with a pair of TALEN DNAs (400ng of each) using Lipofectamine 2000 (Thermo Fisher Scientific). The genomic DNA was purified 48 hours post-transfection. Gene specific primer pairs were used to amplify each of the H2A gene variants. The primer sequences are: H2Aafb3 forward (5′-CAGCAGAAAGCAGCCAAGTGG) and reverse (5′- GCAGGTCAGCCAAGAAGCA); Gm14920 forward (5′ GTACGGTACAAAGGGAG ATG) and reverse (5′-GAGCAGGTCAGCCAAGCAGAG); H2afb2 forward (5′-CAGGTCAGCAGAGAGCAATT) and reverse (5′-CTCCATACTGCTGTAGACCT); and pseudo Lap forward (5′- GTCAGCAGAATGCAGCCAAATAT) and reverse (5′-CAAGCCAGTAGCCAACATCAAG). The PCR products were treated with Surveyor Nuclease (Cel-1) and resolved on PAGE. The nuclease activity of TALENs was measured by quantifying the proportion of the cleaved DNA fragments.

### Talen mRNA synthesis

Plasmids (in Sangamo backbone) were linearized with *XbaI* and the products were purified (PCR Cleanup kit, Qiagen). Using linearized plasmid as template, capped and poly(A)-tailed mRNA transcripts were produced using mMESSAGE mMACHINE T7 Ultra Transcription Kit (Life Technologies) following manufacturer’s instructions. mRNA was purified using a MEGAclear kit (Life Technologies) and was eluted in RNase-free water and single-use aliquots were frozen.

### Production of Talen-mediated genetically modified mice

All mouse work was conducted according to protocols approved and authorised by the University of Queensland Animal and the Australian National University Ethics Committee (Protocol A2014/33). Prior to injection, mRNA was diluted to 10 ng/μ! in filtered RNase-free microinjection buffer (10 mM Tris-HCl, pH 7.4, 0.25 mM EDTA). TALEN mRNA was delivered into the pronucleus of single-cell embryos (FVB), using standard techniques. Injected embryos were cultured overnight to the two-cell stage, then surgically transferred into oviducts pseudopregnant CD1 mice, using standard techniques.

### Analysis of potential founders and genotyping of successive generations

Transgenic founders with mutated H2A.B.3 genes were identified among live born mice by PCR (**Supplementary Table 8**) genotyping of ear notch tissue. Genomic DNA was extracted from ear punches using Quick Extract Solution (Epicentre) following manufacturer’s instructions. The H2afb3, gm14920 and H2afb2 genes were amplified in 25μl volume using the same gene-specific primers as for the Cel1 assay. PCR mix (1x Platinum Taq buffer, 1.5mM Mg2Cl; 0.2mM dNTP; 0.2μM of forward and reverse primer; 2 units of Platinum Taq polymerase (Thermo Fisher)) was amplified using 28 cycles of: 95°C 30s denaturation, followed by 56-65°C, 30s annealing and 72°C, 4min extension.

### Exome Sequencing and bioinformatic analyses

The male mice that were chosen for exome sequencing were genotyped prior to sequencing to confirm that all three generations had the same H2A.B.3 genotype (H2Afb3 (X^Δ5^Y), gm14920 (X^Δ10^Y) and H2Afb2 (X^Δ64^Y)). The purified gDNA was subjected to exome enrichment using the Agilent SureSelect^XT2^ All Exon Kit. The exome capture efficiency was uniformly high with approximately 45-50 *%* of all DNA sequenced being exonic. Sequenced mouse exomes were analyzed using our in-house variant detection pipeline described in detail previously ^20,21^. In short, reads were aligned to the mouse reference genome mm10 using BWA ^22^ and sorted BAM alignment files were generated using SAMTools ^23^. Candidate PCR duplicate reads were annotated using Picard software (http://broadinstitute.github.io/picard/), specifically the MarkDuplicate command. SNVs and small indels were called using SAMTools Mpileup ^24^ and variants were annotated using the Variant Effect Predictor ^25^. To detect larger structural variants, both Pindel ^26^ and Janda (unpublished; https://sourceforge.net/projects/janda/) were utilised. To reduce the number of total variants called, variants specific to the FVBN mouse strain were downloaded from the Sanger Institute ftp site (ftp://ftp-mouse.sanger.ac.uk/REL-1206-FVBNJ/). As these variants were originally reported relative to reference genome mm9, the UCSC liftOver tool (https://genome.ucsc.edu/cgi-bin/hgLiftOver) was used to map the FVBN variants to mm10. These FVBN specific variants were annotated and not considered for further off-target analysis.

For the detection of INDELs in the TALEN mouse exome sequencing data, we followed the GATK recommendations “Best Practices for Germline SNP &; Indel Discovery in Whole Genome and Exome Sequence” ^16^. Briefly, the alignment files in BAM format were realigned and base re-calibration was performed. The cleaned alignments of all sequenced mouse exomes were then used as input for the “HaplotypeCaller” program resulting in raw variants in VCF format, containing both SNVs and INDELs. We then performed hard filtering of the called variants to only include INDELS and applied a quality cut-off score of 100 to include only high-confidence INDEL calls. In order to identify potential TALEN off-targets, we inspected the frequency of INDELs in each generation of TALEN mice, expecting a “dilution” effect due to back-crosses.

In order to search the mm10 reference genome for potential off-target sites, we used the *in silico* PCR function of the UCSC Genome Browser at http://genome.ucsc.edu/cgi-bin/hgPcr ^27^. We used the TALEN target sequences as forward and reverse primers, and relaxed the minimum perfect match parameter to 8 nucleotides (nt) and constrained the maximum product size to 100bp.

### Experimentally validation of putative indels

To interrogate whether the 19 putative indels are present in NM4 G1-G3 mice, the gDNA of wild type FVB/NJArc in-house mice and NM4 G1-G3 KO mice were amplified with primers surrounding the predicted indels, listed in Supplementary Table 9. The PCR products were analyzed by electrophoresis (7% acrylamide gel) and Sanger sequencing.

## ACKNOWLEDGMENTS

We thank Tara Davidson for assistance with microinjections of the TALENs and Dan Andrews for bioinformatics support in analyzing the exome sequencing data. We acknowledge the excellent high-throughput DNA sequencing service provided by our in-house Bimolecular Research Service headed by Stephanie Palmer, and the mouse technicians responsible for the breeding and care of the mice. This work was supported by a NHMRC project grant to D.T. (1058941).

## AUTHOR CONTRIBUTIONS

T.S. and D.T conceived the study and wrote the paper. T.S. devised the analyses. M.F. and S.K. performed exome analyses. L.Z., E.R., and P.G. designed the TALENs and performed the Dual-Luciferase Single Strand Annealing Assay (DLSSA) and the Cel1 cleavage assays. J.B. and P.K. generated the TALEN founder mice. N.D.A. performed the screening, genotyping, Cel1 assays, prepared exome-seq samples and produced the KO mice.

